# A pseudogene of *caffeic acid-o-methyltransferase (COMT)* in *Acacia mangium:* Comparative analysis with other *COMT* plant promoters

**DOI:** 10.1101/269654

**Authors:** Azmah Abdul Latif, Sheh May Tam, Wickneswari Ratnam

## Abstract

*Acacia mangium* is a prominent tree species in the forest plantation industry of Southeast Asia, grown mainly to produce pulp and paper, and to a lesser extent wood chips and solid wood products. Lignin, a natural complex polymer used by plants for structural support and defence, has to be chemically removed during the production of quality paper. Delignification is very expensive and moreover, is an environmental pollutant. Understanding the complex mechanisms that underlie the regulation of lignin biosynthetic genes requires in-depth knowledge of not only the genes involved but also their regulatory elements. Using Thermal Asymmetric Interlaced PCR, a 770 bp promoter sequence with 93% identity with *COMT1* gene from *Acacia auriculiformis* × *A. mangium* hybrid was isolated from *A. mangium*. Bioinformatics analysis revealed the presence of cis acting elements commonly found in other lignin biosynthesis genes such as TATA box, CAAT box, W box, AC-I and AC-11 elements. However, a nonsense mutation that created a premature stop codon was found on the first exon. Modelling of MYB transcription factor binding site on this newly isolated pseudogene shows it has binding sites for important transcription factors involved in lignin biosynthesis both in *Arabidopsis thaliana* and *Eucalyptus grandis*. Given the remarkable structures of its regulatory region, the possible structure of its transcript was detected using Mfold. Results show the transcript are capable of forming stem loop structures, a characteristic commonly attributed to presence of miRNA. Possible functions of pseudo*AmCOMT1* were discussed.

## INTRODUCTION

Lignin is a complex aromatic polymer mainly found in plant secondary cell wall (Boerjan et al. 2003) which plays an important role in plant development (Jones et al. 2001) and defence mechanism (Lauvergeat et al. 2001). Presence of lignin on the secondary cell wall of plants gives its rigidity and upright structure (Jones et al. 2001). Lignin polymer consists of three different monomers, which is p-hydroxyphenyl (H), guaiacyl (G) and syringyl (S) units. Angiosperms lignin consists of mainly G and S subunit. Lignification toolbox in many plant species consists of a set of family protein which functions at different level in the production of lignin monomer. Genome-wide characterisation of Arabidopsis genome found 10 enzymes which functions in lignin biosynthesis pathway (Raes et al. 2003).

Caffeic acid-o-methyltransferase (COMT) is one of the enzyme specifically involved in S unit production (Li et al. 2000). Similar to other genes involved in lignin biosynthesis pathway, *COMT* exist as gene family in many plant species. There are high numbers of genes with high sequence similarity with *COMT* in every plant species. For example, there are 13 *COMT-like* gene in *Arabidopsis thaliana* (Raes et al. 2003) and 25 *COMT* coding sequence in *Populus tricchocarpa* (Shi et al. 2010). However, only one gene was shown to have high expression in xylem tissues and is predicted to have major function in lignification during development (Raes et al. 2003; Shi et al. 2010). *Eucalyptus grandis COMT* gene family was shown to have expanded with seven genes and only one was shown to have high expression in xylem tissues (Carocha et al. 2015). The gene family expansion was determined to be results of tandem duplication and segmental duplication event, followed by functional divergence (Carocha et al. 2015). COMT and other genes which function specifically in S lignin production pathway are predicted to have younger evolutionary origins than genes involved in G lignin production (Peter & Neale 2004).

*Acacia mangium*, an important forest tree species has an estimated worldwide plantation of 1.4 M ha (Griffin et al. 2011). It is planted widely in South East Asia and used mainly for production of pulp and paper (Griffin et al. 2011). Transcriptome sequencing on *Acacia mangium* discovered all the ten monolignol biosynthetic genes involved in lignin biosynthetic pathway (Wong et al. 2011). *COMT* and *CCR-like* gene were also isolated and characterised from *Acacia auriculiformis* × *A. mangium* hybrid (Sukganah et al. 2013).

Regulatory elements involved in the complex regulatory network of lignin biosynthesis include promoter structures, transcription factors and miRNAs (Shi et al. 2010; Grima-Pettenati et al. 2012; Ong & Wickneswari 2011). Genes involved in lignin biosynthesis are known to be co-regulated through binding of MYB transcription factors on their promoter region (Zhao & Dixon 2011). R2R3 MYB transcription factors are also found to be more evolved in woody plants genome (Soler et al. 2015). Through comparative genomic analysis, several sub groups were identified only present in woody plant species and some of its member are later found to also be involved in lignin biosynthesis (Soler et al. 2016).

In this study, a new promoter of a pseudogene with high sequence similarity with COMT is isolated and characterized in silico. The regulatory region of this gene exhibits outstanding characteristics, with presence of ACI and ACII elements, and other cis acting elements commonly found in lignin biosynthetic genes. Comparative analysis with other real *COMT* promoters shows this promoter has the binding site for some important R2R3 MYB transcription factors involved in regulation of lignin and secondary cell wall biosynthesis. The transcript is capable to form stem loop structure. The possible function of this pseudogene is discussed.

## Materials and Method

Young leave samples of *Acacia mangium* were obtained from mature trees grown at plot W, Plant Biotechnology Centre, National University of Malaysia, Bangi, Malaysia. Genomic DNA was extracted from 100mg of fresh leaves using QIAGEN DNeasy Plant Mini Kit (Qiagen, Germany) following manufacturer’s protocol. Isolation of *COMT* promoter sequence was performed using Thermal Asymmetric Interlaced PCR (TAIL-PCR). Two gene specific primers i.e. GSP1 and GSP2 (Sukganah et al. 2013) were used as reverse primers with an arbitrary degenerate primer AD4 (Thanh et al. 2012) as the forward primer (Table 1). TAIL-PCR was adapted from Liu and Whittier (1995) and briefly described below.

**Table 1.** Thermal Cycling Conditions

| Reaction | Number of Cycles | Thermal cycling conditions |
| --- | --- | --- |
| Primary PCR | 1 | 93°C, 1 min; 95°C, 1 min |
|  | 5 | 94°C, 30 s; 62°C, 1 min; 72°C, 2.5 min |
|  | 1 | 94°C, 30 sec; 25°C, 3 min; *72°C, 2.5 min |
|  |  | 94°C, 10 s; 68°C, 1 min; 72°C, 2.5 min |
|  | 15 | 94°C, 10 s; 68°C, 1 min; 72°C, 2.5 min |
|  |  | 94°C, 10 s; 29°C, 1 min; 72°C, 2.5 min |
|  | 1 | 72°C, 5 min |
| Secondary PCR | 12 | 94°C, 10 s; 64°C, 1 min; 72°C, 2.5 min |
|  |  | 94°C, 10 s; 64°C, 1 min; 72°C, 2.5 min |
|  |  | 94°C, 10 s; 29°C, 1 min; 72°C, 2.5 min |
|  | 1 | 72°C, 5 min |
| Tertiary PCR | 20 | 94°C, 15 s; 29°C, 30 s; 72°C, 2 min |
|  | 1 | 72°C, 5 min |

\* ramping to 72°C, over 3 min

The primary PCR reaction mixture consisted of 1 U of HS Taq polymerase (Takara Bio Inc., Japan), 1× Taq polymerase buffer, 200 μM dNTPs, 0.2 μM GSP1 primer, 5 μM of the AD primer, and 50 ng of genomic DNA. The secondary PCR reaction consisted of 1× Taq polymerase buffer, 200 μM dNTPs, 0.8 U of HS Taq polymerase, 0.2 μM GSP2 primer, 4 μM of the AD primer used in the primary reaction, and 50 fold dilution of the primary PCR product. The tertiary PCR mixture consisted of 1× Taq polymerase buffer supplied with the enzyme, 200 μM dNTPs, 0.5 U of HS Taq polymerase, 3 μM of the AD primer used in the previous reactions, 0.3 μM GSP2 primer, and 10 fold dilution of the secondary PCR product. The thermal cycling conditions are shown in Table 2. PCR products were visualized by electrophoresis on 1.0% (w/v) agarose gels, purified using NucleoSpin Gel and PCR Clean-up kit and sequenced.

**Table 2.**
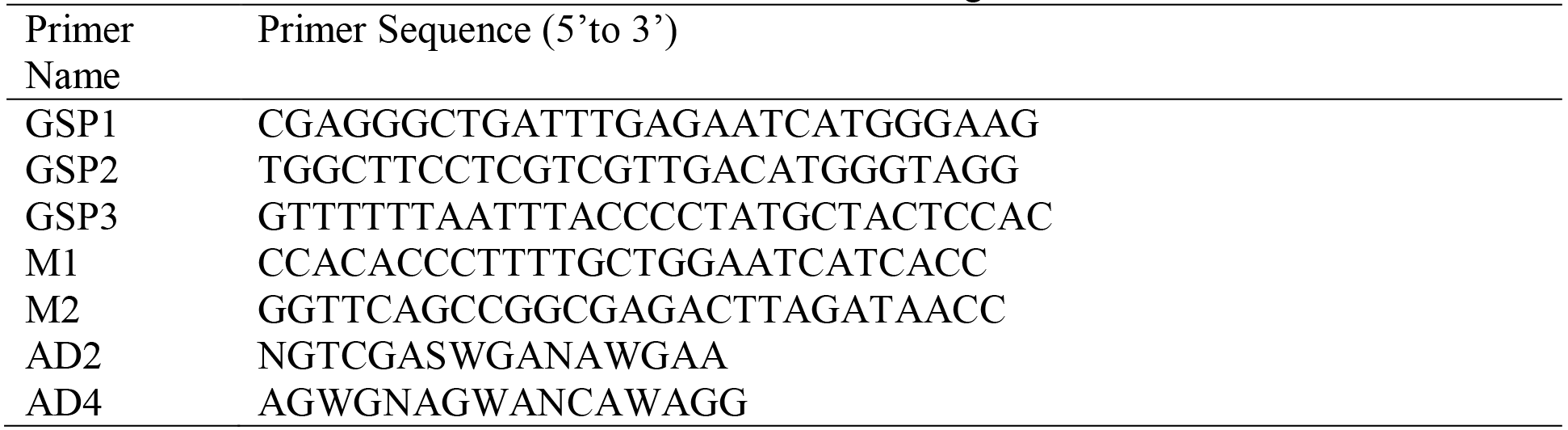
Primer utilized for further isolation of 5’ and 3’ region

After the first round of PCR and sequencing, new primers (GSP3, M1 and M2) were designed based on the sequence of the first fragment obtained to further amplify the 5’ and 3’ regions (Table 2). GSP1, GSP2 and GSP3 were used in combination with AD2 to amplify the promoter region. Primers M1 and M2 were used to further amplify the 3’ region. All PCR products were visualized by electrophoresis on 1.0% (w/v) agarose gels. Strong, clear bands were extracted and purified using the NucleoSpin Gel and PCR Clean-up kit (Macherey Nagel) before being sent for sequencing (First Base).

### Sequence Analysis

Sequences were manually cleaned and assembled using BioEdit (Hall 1999) before being subjected to similarity search using BLASTn (Altschul et al., 1990). The promoter sequence was aligned to known *COMT1* promoter sequence (HQ317735.1) from *Acacia auriculiformis* × *A. mangium* hybrid (Sukganah et al. 2013) using Clustal Omega (Sievers et al. 2011). Putative *cis*-acting elements were identified using PlantCARE (Lescot et al. 2002), while the ORF of the gene was detected using ExPASy Translate Tool. The secondary structure of the CDS sequence was predicted using Mfold, a tool for predicting the secondary structure of RNA and DNA (Zuker 2003).

PlantTFDB is a plant transcription factor database which allows the prediction of transcription factor binding on promoter sequence (Jin et al. 2017). Transcription factor binding site on pseudo*AmCOMT1*, *AtCOMT* and *EgrCOMT1* were predicted using PlantTFDB (Jin et al. 2017). R2R3 MYB transcription factor binding site on pseudo*AmCOMT1* and *AhgCOMT1* were further modelled with *AtCOMT* and *EgrCOMT1* promoters using Cytoscape (Shannon et al. 2003). All software was used on default parameter.

## RESULTS AND DISCUSSIONS

### Isolation and analysis of Promoter

The initial TAIL-PCR successfully amplified a 400bp fragment of DNA from *A. mangium*. Analysis of the DNA sequence of this fragment showed part of the promoter, associated 5’UTR and exon 1 of the *COMT* gene (270bp), with 93% similarity with *AhgCOMT1* from *Acacia auriculiformis* × *Acacia mangium* hybrid. However, a nonsense mutation was detected on exon 1, and in order to confirm this and to increase the length of promoter obtained, new TAIL-PCR primers were successfully used to amplify and extend the 5’ and 3’ regions of this DNA fragment. This resulted in a 800bp fragment with 92% similarity to *AhgCOMT1*. Regions of high sequence similarities are detected within 211bp of the promoter region to ~110bp after the start (ATG) codon (Figure 1). Since the presence of the nonsense mutation was confirmed, this sequence was thus named pseudo*AmCOMT1* (Genbank accession number: MF488717).

The length of the promoter of pseudo*AmCOMT1* is 697 bp and PLANTCARE analysis of the promoter sequence shows presence of multiple *cis*-acting elements (Figure 1). The core promoter element, TATA box was found at position −22. The 5UTR Py-rich stretch, a *cis*-acting element which confers high transcription levels was identified at −261. The elements ACI and ACII which are commonly found in genes involved in lignin and phenylpropanoid pathway, were identified at position −77 and −40 respectively. AC elements are important regulatory elements commonly found in lignin biosynthetic genes and known to be the target for binding of both activator and repressor protein. Presence of AC elements in almost all promoters of lignin biosynthetic genes enables coordinated gene expression towards xylem tissues (Zhou et al. 2009). AC elements were previously reported to be absent in most *COMT* genes (Raes et al. 2003), but expected to be more degenerative and could not be picked up by bioinformatics analysis (Zhou et al. 2009).

**Fig. 1.**
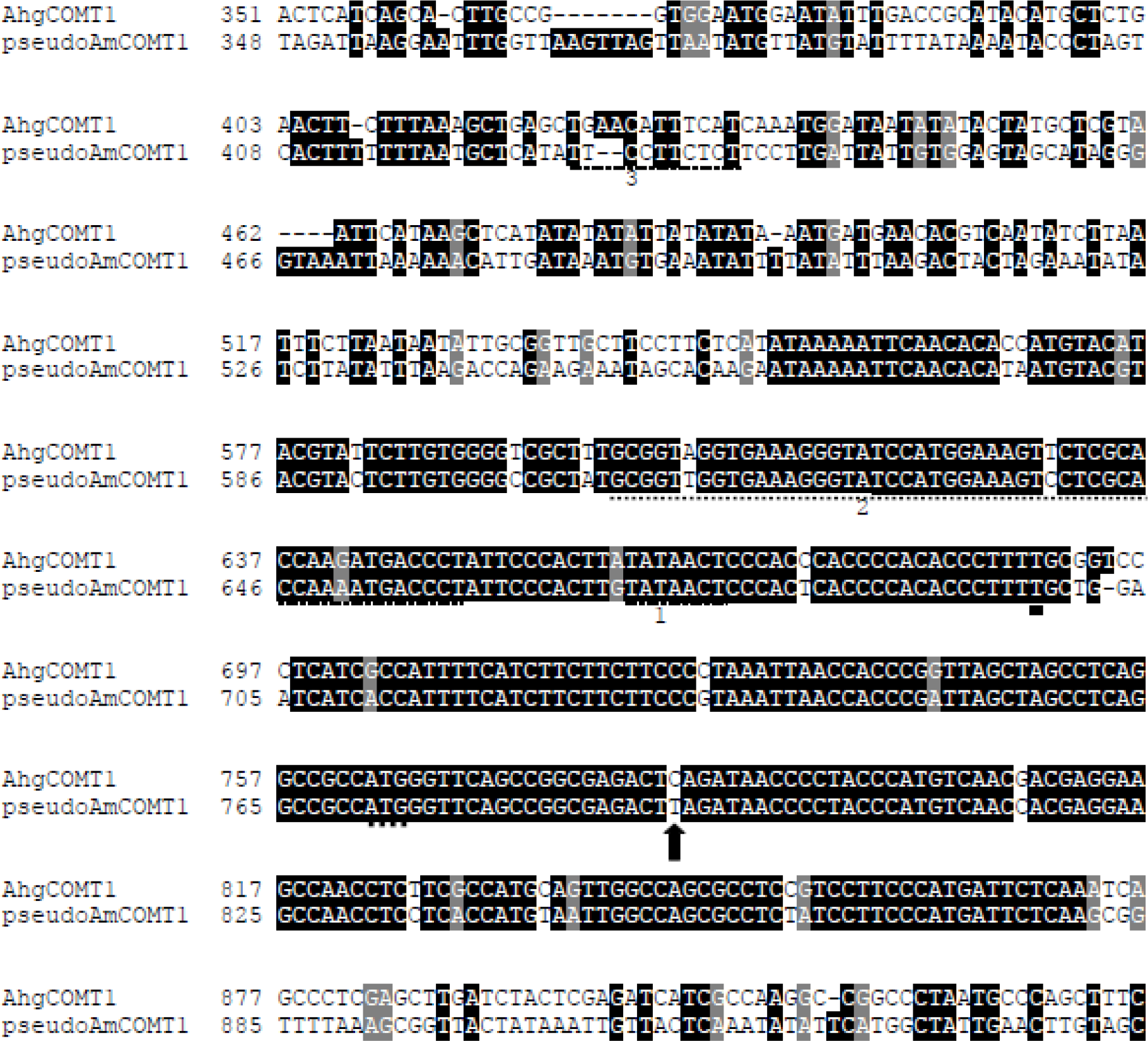
Alignment of *pseudoAmCOMT1* and *AhgCOMT1*. Transcription start site is marked with thick underline. Translation start site (ATG) is marked with dashes. Important *cis* acting elements are underlined with dashes and numbered. 1: TATA box, 2: ACI and ACII elements, 3: 5UTR Py-rich stretch. Nonsense mutation that created premature stop codon marked with an arrow.

Multiple *cis* elements that function in light responsiveness such as the ATCT-motif, Box I, GA-motif and GT1-motif were also identified in this promoter at positions −345, −256 and −330 respectively. This is not surprising as expressions of lignin genes are also known to be affected by light, the circadian clock and sugar levels (Rogers et al. 2005). Other elements identified include W box which functions in plant defence against pathogens and two motifs GCN4 and Skn-1 which regulate endospermic gene expression, at −619 and −43. Abscisic acid responsiveness motif, ABRE was found at −110 whilst an ARE motif that functions in anaerobic induction was shown at −496. P-box, a motif implicated in gibberellin response was found at +2. Overall, it is noted that similar sets of *cis*-acting elements were also reported from promoters of other lignin genes such as *cinnamoyl-CoA reductase (CCR)* and *cinnamyl alcohol dehydrogenase (CAD)* (Prashant et al. 2012; Lacombe et al. 2000).

### Transcription Factor Binding Site Comparative Analysis

*In silico* analysis of pseudo*AmCOMT1* revealed the presence of multiple cis-elements that may serve as binding sites for transcription factors with important functions in lignin biosynthesis. To further explore this, PlantTFDB (Jin et al. 2017) was used to obtain comparative models of transcription factors binding on regulatory regions of pseudo*AmCOMT1*, *AhgCOMT1* and *AtCOMT* (Figure 2) as well as pseudo*AmCOMT1*, *AhgCOMT1* and *EgrCOMT1* (Figure 3). The model shows interactions with MYB gene family members, such as AtMYB46 and AtMYB83 which play the role of master regulatory switch in secondary cell wall biosynthesis (Zhong and Ye 2012; McCarthy et al. 2009; Ko et al. 2009). Both AtMYB46 and AtMYB83 function to activate AtMYB58, AtMYB63 and AtMYB85 which are on the next level of the regulation cascade (Zhong et al. 2007, McCarthy et al. 2009). However, it was recently shown that their mode of action was not restricted to just downstream activation of another set of transcription factors, but instead they could also directly activate some lignin genes through secondary wall MYB-responsive element (SMRE) binding site (consensus motif ACC(A/T)A(A/C)(T/C)) in the promoter region (Zhong & Ye 2012).

**Fig. 2.**
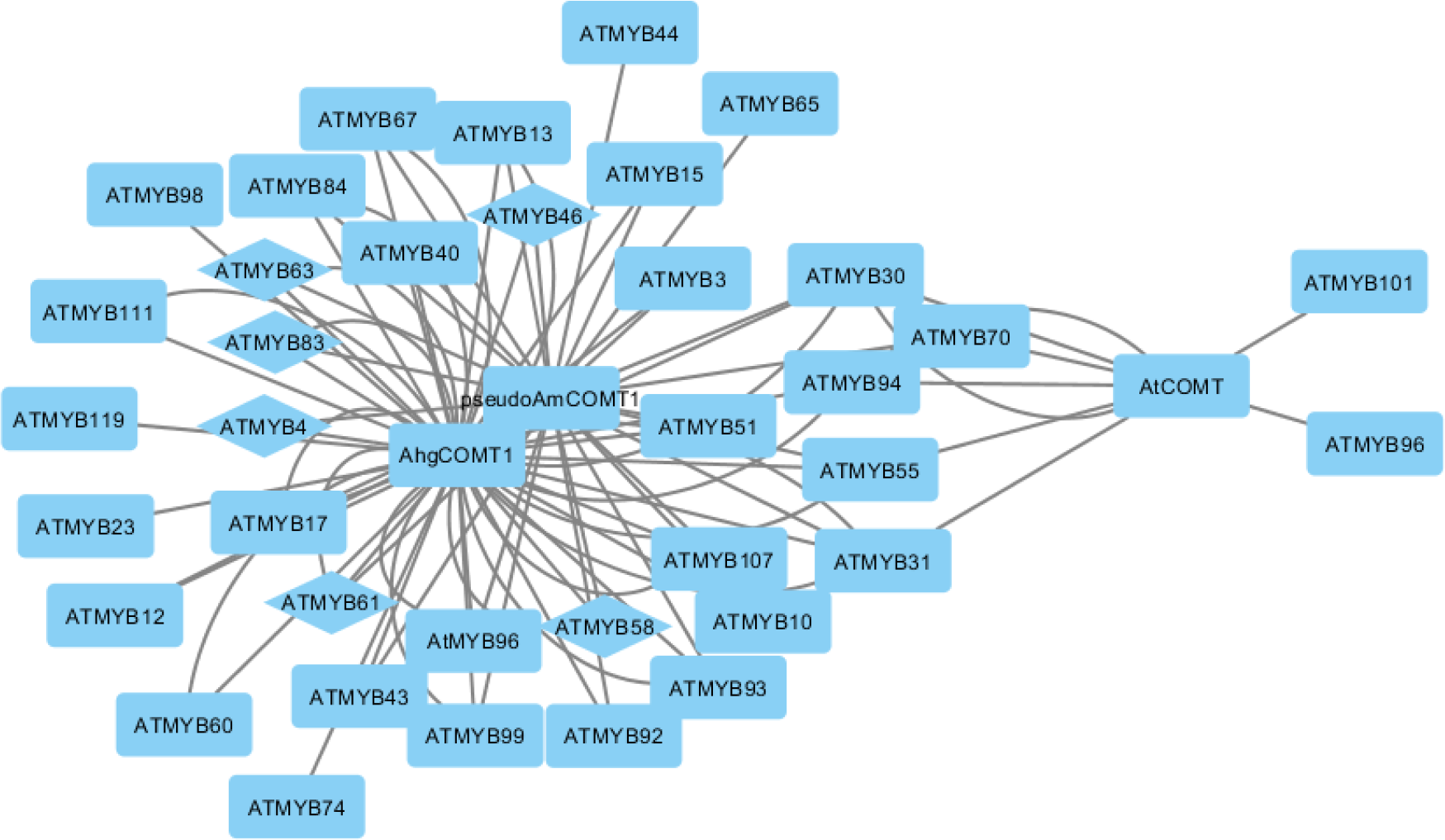
Transcription factor binding site model of pseudo*AmCOMT1*, *AhgCOMT1* and *AtCOMT*. Important transcription factors involved in lignin and secondary cell wall biosynthesis are shown in diamond shape.

AtMYB58, AtMYB63 and AtMYB85 are three transcription factors which function specifically in lignin biosynthesis regulation. Pseudo*AmCOMT1* and *AhgCOMT1* are shown to have binding sites for both AtMYB58 and AtMYB63. These three transcription factors were found to coordinately activate the whole lignin biosynthetic genes by binding to the AC elements in their promoter region, except for *F5H* (Zhou et al. 2009). Although the binding sites for MYB58 and MYB63 on *AtCOMT* were not detected using bioinformatics analysis, these two transcription factors were shown to regulate the expression of *AtCOMT* gene (Zhao & Dixon 2009).

**Fig. 3.**
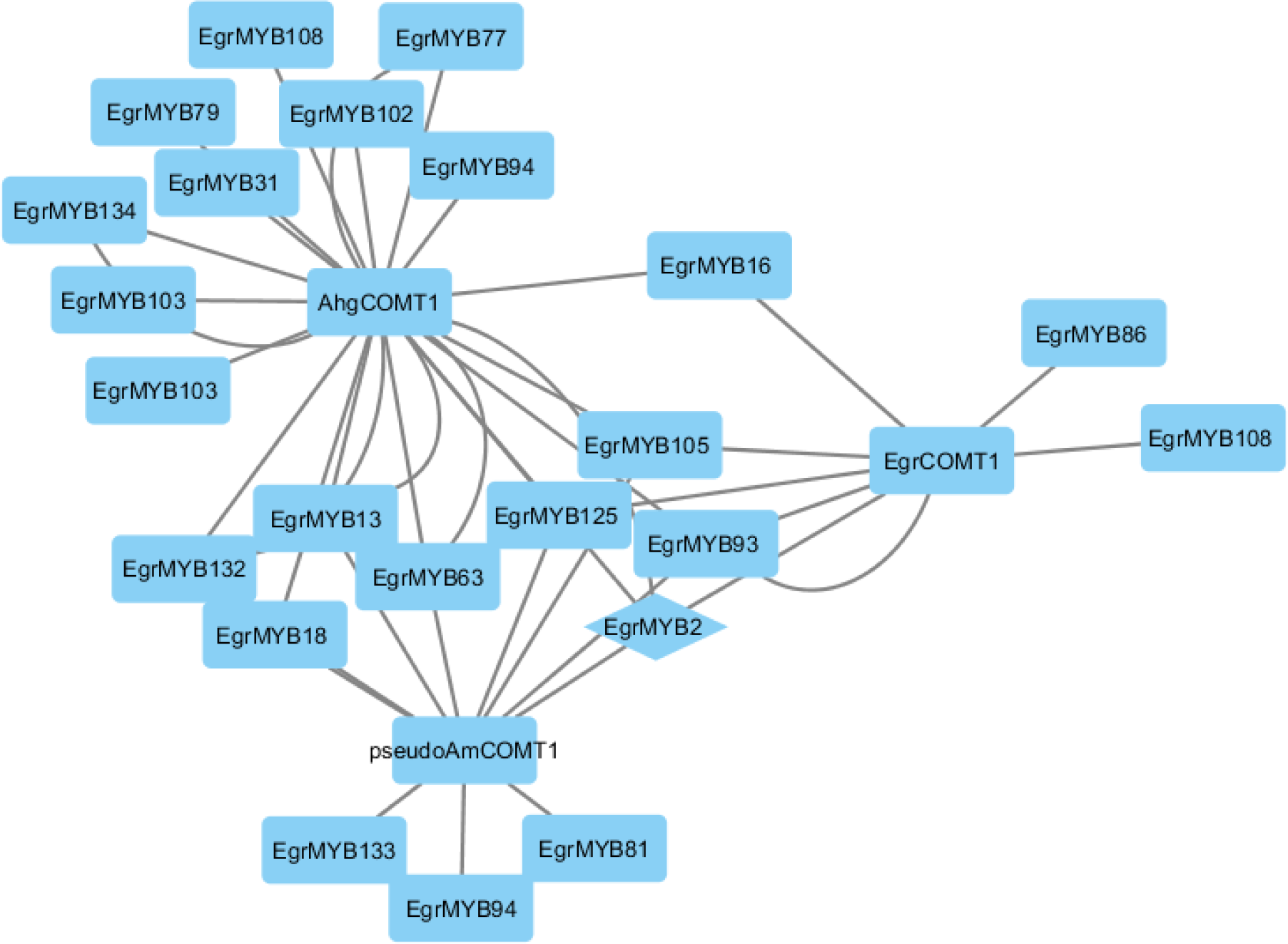
Transcription factor binding site model of pseudo*AmCOMT1*, *AhgCOMT1* and *EgrCOMT1*. Important transcription factors involved in lignin and secondary cell wall biosynthesis are shown in diamond shape.

Pseudo*AmCOMT1* and *AhgCOMT1* are also identified to have binding site for AtMYB61, which was responsible for causing ectopic lignification in *Arabidopsis* when overexpressed (Newman et al. 2004). AtMYB13, AtMYB15 and AtMYB10 which are part of subgroup S2&3 are also predicted to have binding sites on pseudo*AmCOMT1* and *AhgCOMT1* (Soler et al. 2015). Both pseudo*AmCOMT1* and *AtCOMT* promoters are shown to have binding site for AtMYB55 which is a member of subgroup S13 (Soler et al. 2015). Members of both subgroups (S2&3 and S13) are predicted to be involved in secondary cell wall formation in *Arabidopsis*. In addition, *AtCOMT* is also shown to have binding sites for AtMYB70, AtMYB31, AtMYB96, AtMYB30, AtMYB101 and AtMYB94. AtMYB96 and AtMYB94 are members of Subgroup 1 which is predicted to be involved in plant defence mechanism (Soler et al. 2015). Pseudo*AmCOMT1* and *AhgCOMT1* are also shown to have binding site for AtMYB4, an important repressor in lignin biosynthesis pathway (Jin et al. 2000).

Results of the transcription factor binding site model for pseudo*AmCOMT1*, *AhgCOMT1* and *EgrCOMT1* predicted all three promoters to have a binding site for EgrMYB2, an important transcription factor known to be a master regulator of secondary cell wall biosynthesis in *Eucalyptus* (Goicoechea et al. 2005). Overexpression of *EgMYB2* in tobacco caused dramatic increase in secondary wall thickness, as well as alteration in lignin profiles, where the S:G ratio was increased. Transcript analysis showed upregulation of *EgMYB2* caused an increase of 40 folds of *COMT*, *CCoAOMT*, *F5H* and *C3H* genes expression while *CCR* and *CAD* were upregulated five fold (Goicoechea et al. 2005). This model indicates the difference might be partly because of the genes’ promoter structure. Both pseudo*AmCOMT1* and *EgrCOMT1* promoters also have binding site for EgrMYB93 with the former with the former having additional binding sites for EgrMYB132 and EgrMYB133. All these three transcription factors belong to groups which are expected to have important functions in the regulation of secondary cell wall biosynthesis (Soler et al. 2015). Interestingly, the binding site for EgrMYB81, a member of the ‘woody-expanded subgroup’, S6 (Soler et al. 2015) was detected from pseudo*AmCOMT1*. Members of this subgroup were identified to be present significantly more in woody plant species (Soler et al. 2015).

Given the strong regulatory elements present in the promoter region, the transcript sequence of pseudo*AmCOMT1* was subjected to secondary structure prediction. Results show that the transcript is able to form hairpin structures with ΔG = −1.6 (a) and −1.3 (b & c) (Figure 5). Generally, the ability to form hairpin precursor indicates the formation of miRNA (Ambros et al. 2003) and suggests the possibility of pseudo*AmCOMT1* in regulating its paralogous genes.

**Fig. 4.**
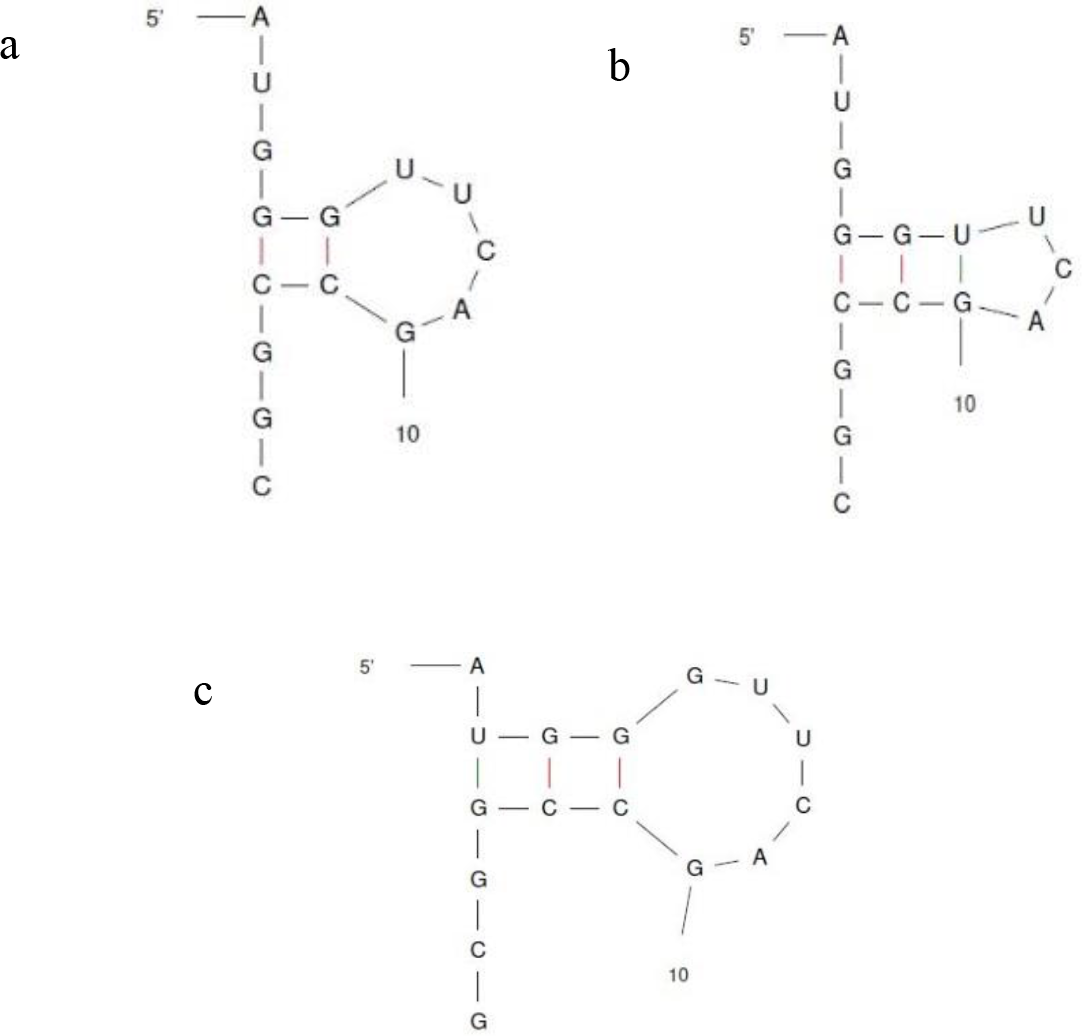
Hairpin structures of pseudo*AmCOMT1* transcript.

*COMT* gene family were identified to be among the most expanded gene family among genes involved in lignin biosynthesis, and biotic defence were predicted to be the driving force for the rapid expansion (Xu et al. 2009). Rate of tandem duplication of *COMT* genes in *Eucalyptus grandis* exceeds overall frequency of tandem duplication within the genome (Carocha et al. 2014; Myburg et al. 2014). Expressions of most of these tandem duplicated genes was found to be strongly responsive towards environmental stimuli (Carocha et al. 2014).

Recent findings shows a group of small interfering RNAs with young evolutionary origins, phasiRNAs targets genes in the same family from where they are derived (Zheng et al. 2015). This group of small interfering RNAs were found to have nearly perfect match to the genes in the same family as their originating loci and play important role in disease resistant within dicot species (Zheng et al. 2015). Some phasiRNAs were found to negatively regulate disease resistance genes in *A. thaliana* (Boccara et al. 2014). Members of COMT gene family, along with other genes involved in lignification, are known to also take part in plant resistance against pathogen (Toquin et al. 2003; Lauvergeat et al. 2001). In fact, resistance against pathogen are expected to be the major driving force of *COMT* gene family expansion (Xu et al. 2009).

Interestingly, downregulation of *COMT* in Poplar and tobacco causes only minor effect on plant morphology. While downregulation of most of the lignin biosynthetic genes causes reduction in lignin content and an abnormal phenotype, downregulation of *COMT* in both cases caused reduction in S lignin unit, but the overall lignin remain unchanged as 5-hydroxyconiferyl alcohol was incorporated (Atanassova et al. 1995; Doorsselaere et al. 1995). This new type of lignin composition was also found to have more cross link and thus, more resistant to degradation (Jouanin et al. 2000). Possible role of pseudo*AmCOMT1* in regulating its orthologous genes however, could only be confirmed through future functional studies.

## ACKNOWLEDGEMENTS

We would like to thank Ministry of Science and Technology and Innovation (MOSTI), Malaysia (02-02-18-SF-1117/ST-2013-012) for funding.

## COMPLIANCE WITH ETHICAL STANDARDS

Conflict of Interest: All authors declare that they have no conflict of interest.

